# Structure of the rabies virus glycoprotein trimer bound to a pre-fusion specific neutralizing antibody

**DOI:** 10.1101/2021.11.16.468852

**Authors:** Heather M. Callaway, Dawid Zyla, Kathryn M. Hastie, Ruben Diaz Avalos, Alyssa Agarwal, Hervé Bourhy, Davide Corti, Erica Ollmann Saphire

## Abstract

Rabies infection is nearly 100% lethal if untreated and kills over 50,000 people annually, many of them children. Existing rabies vaccines target the rabies virus glycoprotein (RABV-G) but generate short-lived immune responses, likely because the protein is heterogeneous under physiological conditions. Here, we report the 3.39Å cryo-EM structure of trimeric, pre-fusion RABV-G complexed with RVA122, a potently neutralizing human antibody. RVA122 binds to a quaternary epitope at the top of RABV-G, bridging domains and stabilizing RABV-G protomers in a prefusion state. RABV-G trimerization involves side-to-side interactions between the central α-helix and adjacent loops, rather than contacts between central helices, and interactions among the fusion loops at the glycoprotein base. These results provide a basis to develop improved rabies vaccines based on RABV-G stabilized in the prefusion conformation.

**One sentence summary:** We report the structure and trimeric interface of pre-fusion rabies virus glycoprotein bound to the neutralizing antibody RVA122.

## Main Text

Untreated rabies infections are nearly 100% fatal, causing 50,000-60,000 human deaths annually and also significantly impacting animal populations (*1*). Rabies vaccines for human and canine use consist of inactivated virus and have existed since the late 1800s, but elicit only short-lived immunity. In humans, levels of vaccine-elicited neutralizing antibodies usually wane 1-5 years after vaccination (*2*) and frequent revaccination is required to maintain protection (*3*). Unvaccinated individuals who are exposed to rabies are given post-exposure prophylaxis consisting of polyclonal antibodies derived from sera of vaccinated individuals or immunized horses and multiple doses of the rabies vaccine (*4*). However, both frequent vaccination and post-exposure treatment are unaffordable in low-income countries where most rabies deaths occur.

Rabies virus is a member of the family *Rhabdoviridae* and the genus *Lyssavirus*. Lyssaviruses are further divided into 3 phylogroups, with phylogroup I including all rabies virus strains and phylogroups II and III containing more distantly related species (*5*). All known lyssaviruses are believed to cause viral encephalitis and the same clinical symptoms as rabies infection. The rabies virus glycoprotein (RABV-G) is the only surface-exposed protein on the virus and is the target of vaccine-elicited neutralizing antibodies. RABV-G shares 57-78% protein sequence identity with other lyssaviruses, but only ∼20% identity with other rhabdoviruses. Pan-lyssavirus antibodies that recognize conserved glycoprotein epitopes are desirable for developing more broadly effective therapeutics, but thus far only a small number have been described in detail (*6*–*9*).

On the viral surface, RABV-G is structurally heterogeneous and only a portion is recognizably trimeric (*10*–*12*). As a class III viral fusion protein, RABV-G undergoes reversible, largely pH-dependent transitions between pre- and post-fusion conformations (*10, 13, 14*). In the pre-fusion conformation, the fusion loops point towards the viral membrane (*7, 12*). Exposure to acidic pH triggers a conformational change to the post-fusion state in which RABV-G elongates and the fusion loops point away from the viral membrane and toward the target cell membrane (*7, 12*). Virions display both pre- and post-fusion conformations over a range of physiological pHs (*13, 15, 16*). This structural heterogeneity may affect the generation of neutralizing antibodies that often target quaternary epitopes and likely contributes to the poor vaccine longevity.

Here, we report the structure of trimeric, wild-type RABV-G bound to the human antibody RVA122, which was isolated from a vaccinated individual as part of an effort to develop neutralizing, pan-lyssavirus antibodies for improved post-exposure prophylactic therapy (*8*). RVA122 is specific for the RABV-G pre-fusion conformation and potently neutralizes multiple phylogroup I lyssaviruses, including rabies (*8*). We show that the RABV-G trimeric interface involves interactions between the central α-helix and adjacent loops and demonstrate the role of the fusion loops in trimerization and stabilization of the pre-fusion conformation. The pre-fusion trimer structure, elucidation of a potently neutralizing antibody epitope, and illumination of the fusion loop structure and activity detailed here can be used to improve vaccines and identify therapeutic drug targets.

### Structure determination

We used cryo-electron microscopy to resolve the structure of the trimeric RABV-G ectodomain (PV strain) in complex with three RVA122 antigen binding fragments (Fabs) to 3.39Å (Fig. 1 and S1, Table S1). RABV-G fusion loops anchor to either cellular membranes that co-purify with the protein or detergent micelles added during grid preparation (Fig. 1A-C). RABV-G trimers with unanchored fusion loops were also present on grids (Fig. 1C), but the fusion loops were too flexible to reach high resolution. Anchored fusion loops also adopted multiple conformations (Movie S1).

**Fig. 1.**
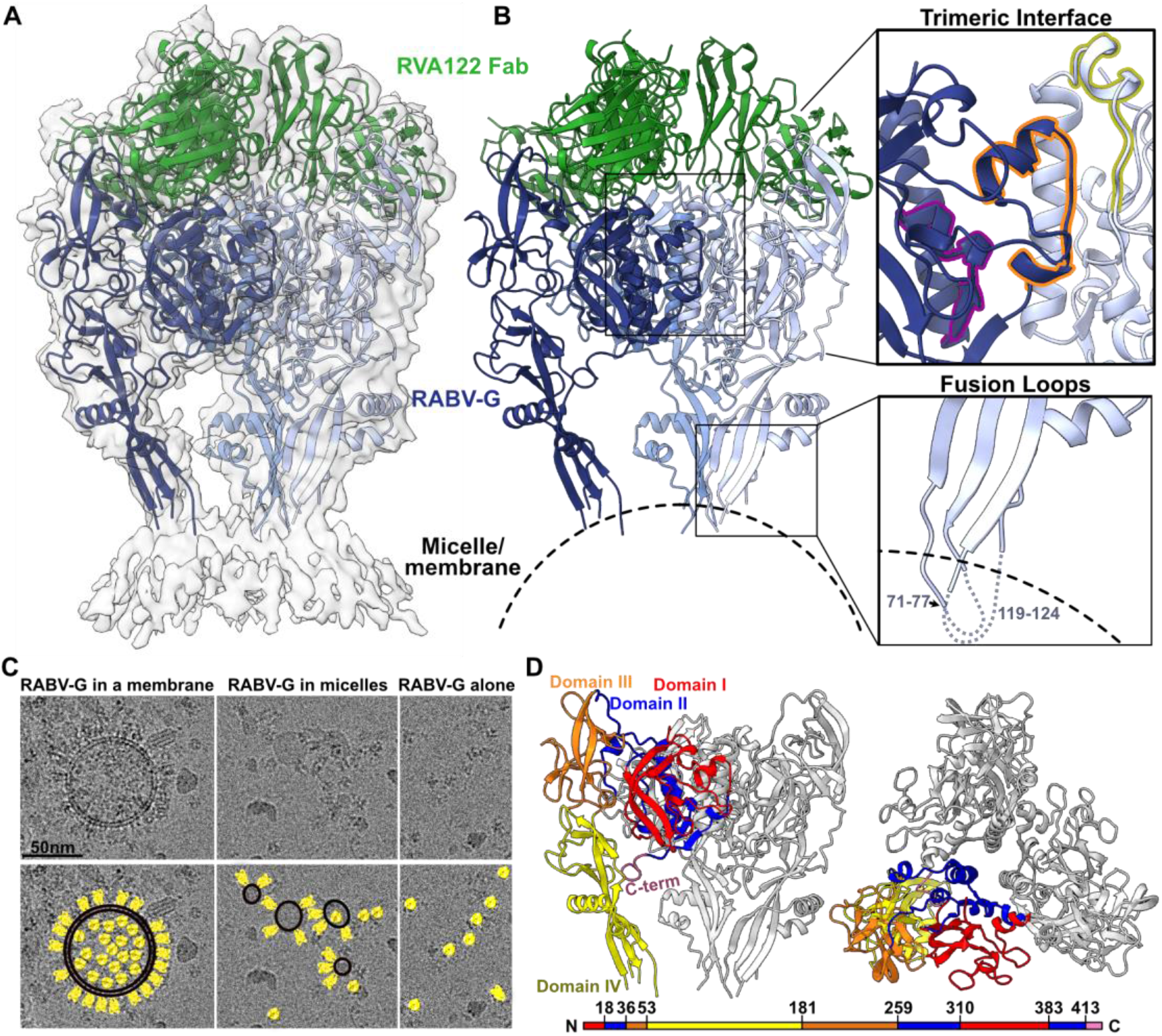
Structure of the pre-fusion RABV-G trimer bound to neutralizing antibody RVA122. **(A)** Molecular model of RABV-G (blue) and RVA122 Fab (green) complex fitted into the corresponding cryo-EM density map. **(B)** RABV-G trimer with magnified views of the trimeric interface (top right) and fusion loops (bottom right). In the trimeric interface, the small helix (purple), bracket loop (orange), and corkscrew loop (yellow) are highlighted. **(C)** (Top) Raw micrographs of glycoprotein complexes embedded in cellular membranes (left), embedded in micelles (center), or unembedded (right). (Bottom) RABV-G (yellow), membranes (black, double outline) and micelles (black, single outline) are indicated. **(D)** Side view (left) and top view (right) of the RABV-G trimer, with Domains I-IV and the C-terminus indicated for one protomer. Colors correspond to RABV-G domains I-IV, also shown in the schematic diagram of the RABV-G sequence.

RABV-G protomers have four linked domains (Fig. 1D) that reposition via hinge regions during the pre-fusion to post-fusion transition (*7*). Domains I and III comprise the solvent-exposed upper half of RABV-G and contain multiple antigenic sites (including the RVA122 epitope) and receptor binding sites (Fig. 1D, S2) (*8, 17*–*19*). Domain II contains a central helix that elongates in the post-fusion transition (*7*), and Domain IV contains the two fusion loops. Domains I and II are also known as the central domain (CD), Domain III as the Pleckstrin homology domain (PHD), and Domain IV as the fusion domain (FD).

### Trimeric interface

Although an α-helix from each RABV-G protomer forms a 3-fold axis of symmetry at the center of the glycoprotein (Fig. 1D), the helices are too distant to interact (Fig. 2A). Instead, the RABV-G trimeric interface primarily consists of lateral interactions among protomers (Fig. 2A and C). Key to these interactions is a loop in domain I that forms a bracket between two short helices (residues 378-384). This bracket loop extends out from one protomer to interact with two sections of the neighboring protomer: the central helix (residues 274-293) and a loop that extends from the central helix to form a corkscrew (residues 259-271) (Fig. 2C). The bracket loop, corkscrew loop, and central helix together form a network of hydrophilic and hydrophobic interactions involving both the main-chain and amino acid side chains (Fig. 2C). The remaining protomer-protomer contacts occur between a small α-helix extending from the bottom of the central helix on one protomer and the central helix of the adjacent protomer (Fig. 2C).

**Fig. 2.**
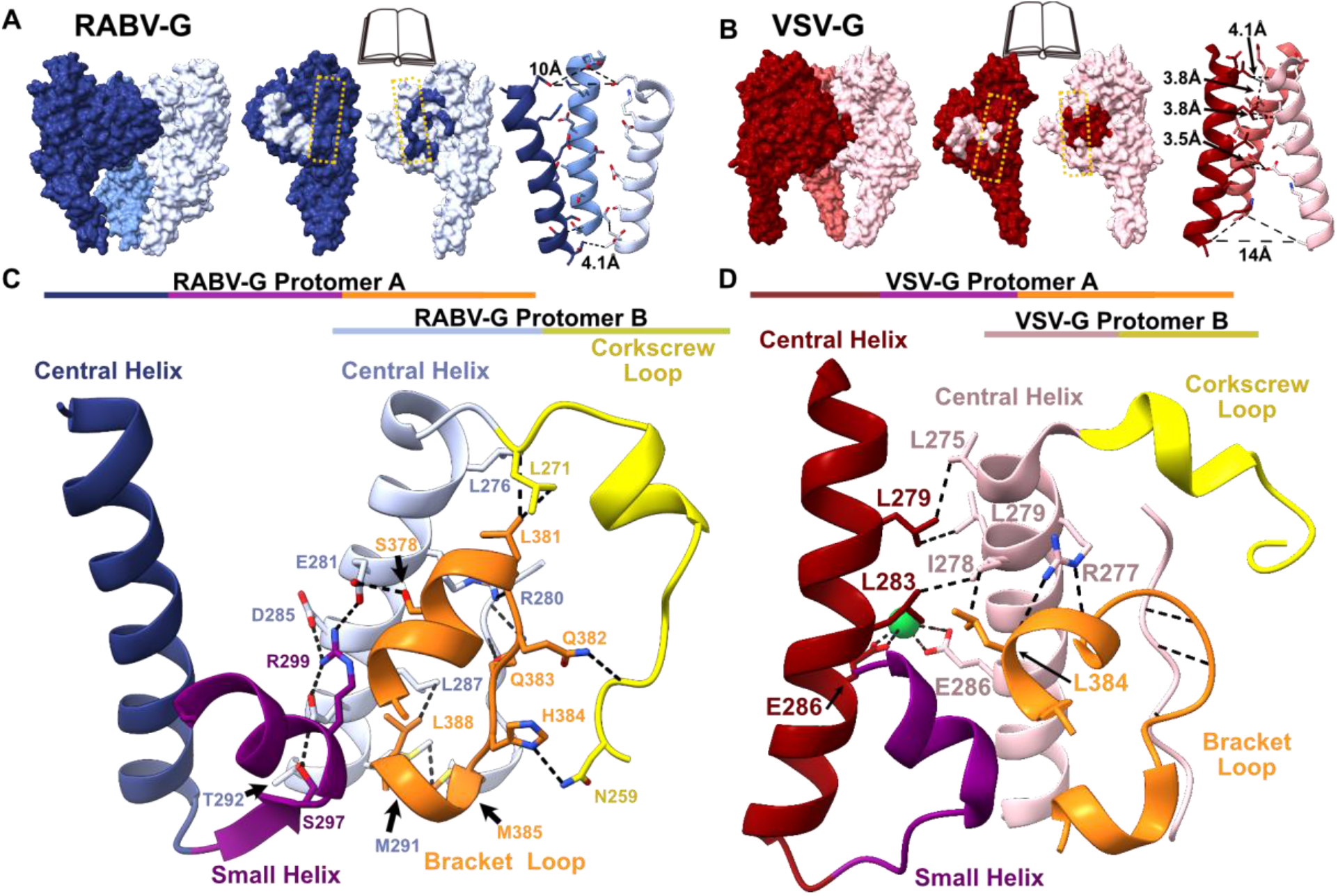
The trimeric interface of pre-fusion RABV-G. **(A and B)** Book-end models of the interface between protomers of RABV-G (A) and VSV-G (B) (PDB ID: 5i2s) with corresponding contacts from adjacent subunits shaded (left and center). The yellow dashed line indicates the central helix. On the right, the RABV-G and VSV-G central helices and distances between side chains are shown. **(C and D)** Magnified view of the RABV-G (C) and VSV-G (D) trimeric interface, with the corkscrew loop (yellow), the bracket loop (orange), and the small helix (purple). Dashed black lines denote hydrogen bonds and hydrophobic interactions.

In this structure, the bracket loop and C-terminus adopt significantly different conformations compared to a previously reported structure of monomeric RABV-G, in which the corkscrew loop is unresolved (Fig. S3) (*7*). In the monomeric structure, residues 373-389 of the bracket loop instead form a helix, which extends into a long, flexible loop that projects sideways and makes crystal contacts with another monomer (Fig. S3) (*7*). This extended conformation disrupts the trimeric interface. The alternative conformations of the bracket and corkscrew loops suggest that the trimeric interface is unstable, which could explain why the trimeric RABV-G structure has been so elusive.

To better understand the trimeric interface, we examined vesicular stomatitis virus (VSV), another member of the rhabdovirus family and the most closely related virus to rabies for which a trimeric glycoprotein structure is available (*20, 21*). The overall organization of RABV-G and VSV-G is similar, but the relatively low sequence identity (∼22%) leads to differences in interactions at the trimeric interface. In RABV-G, the central helices are arranged as a cone with a wide top and narrow bottom (Fig. 2A). In contrast, in VSV-G, the central helices form a cone with a narrow top and wide bottom (Fig. 2B), and the glycoprotein assembles primarily through interactions between the central helices, with the bracket loop shifted and forming backbone-backbone hydrogen bonds with the adjacent protomer (Fig. 2D).

Differences in residue composition between RABV-G and VSV-G also influence propensity to trimerize and trimer stability. During the transition to the post-fusion conformation, the VSV-G central helices elongate at the top, introducing additional contact points between protomers (*21*). As a result, VSV-G trimer stability increases with decreasing pH, which favors the transition to the post-fusion conformation (*21, 22*). In RABV-G, where the pre-fusion central helices are arranged into a cone with a wide top, central helix elongation in the post-fusion transition would not be expected to yield strong inter-protomer contacts. Indeed, RABV-G ectodomain trimers destabilize with decreasing pH, as evidenced by a four-fold lower inter-monomer binding affinity at pH 5.5 than at 7.4 (Fig. S4). Under cryo-EM conditions at pH 5.5, our RABV-G ectodomain has a monomeric and heterogeneous conformation, and post-fusion RABV-G, unlike VSV-G, was crystallized as a monomer (*7*). These findings support our structural evidence that the pre-fusion trimeric interface of RABV-G is built not primarily by a central core of tightly interacting helices and hydrophobic contacts, but instead by lateral interactions between the central helix and adjacent loops.

### Neutralizing antibody recognition

Challenges in developing longer lasting rabies vaccines include the lack of the quality and durability of the resulting neutralizing antibody response, and the lack of efficacy of most antibodies against other emerging and circulating lyssaviruses [22,23]. Monoclonal antibody RVA122 is the type of antibody desired from rabies immunization. RVA122 potently neutralizes rabies virus with an IC_90_ of ∼0.1ng/mL and neutralizes multiple other phylogroup I lyssaviruses with similar efficacy (*8*). Three copies of RVA122 bind to the top of the RABV-G trimer, with each Fab adhering to a quaternary epitope that bridges domains I and III and contacts domain II on an adjacent protomer (Fig. 3). This footprint is unique compared to the two other RABV-G binding antibodies with known structures: RVA20 and 523-11. Both RVA20 and 523-11 neutralize rabies virus, but each binds to a single domain that has the same conformation in pre- and post-fusion RABV-G (Fig. 3A) (*6, 7*).

**Fig. 3.**
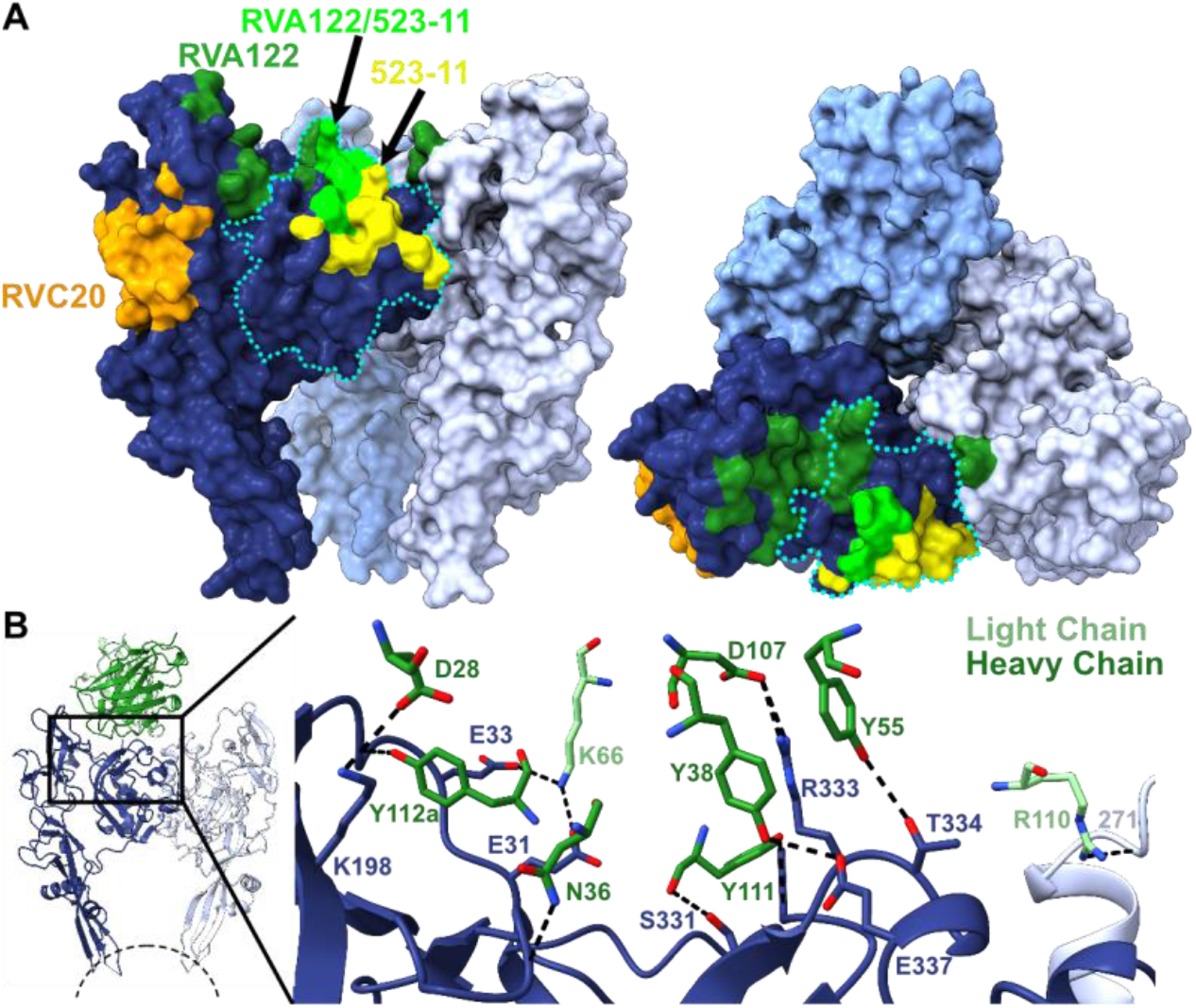
RVA122 binds a quaternary epitope. **(A)** Binding footprints for antibodies RVA122 (dark green), 523-11 (yellow) and RVA20 (orange), and overlap in footprints between RVA122 and 523-11 (bright green). RABV-G Domain I is outlined in a teal dotted line. **(B)** Interactions between RVA122 (green) and RABV-G (dark and light blue). RVA122 residues are labeled in IMGT format.

RVA122 binding increases the proportion of RABV-G trimers visible via cryo-electron microscopy by over 30-fold, making high-resolution reconstruction possible, and also locks RABV-G into the pre-fusion conformation. RVA122 likely neutralizes rabies virus by inhibiting the transition to the post-fusion conformation, as the antibody remains bound below pH 5 after negative staining. Residue contacts between RVA122 and RABV-G include domain I residues S331, R333, T334, and E337 and domain III residues E31, E33, and K198, most of which are highly conserved among type I lyssaviruses (Fig. S7), explaining why RVA122 is broadly neutralizing. RVA122 light chain residue R110 forms a single contact with domain II residue L271 on the neighboring protomer (Fig. 3B). Mutation of R110 to Ala or Glu did not significantly impact binding affinity; this contact therefore does not appear to be critical (Fig. S8). Because mutation of RVA122 residue R110 has a negligible effect on antibody binding, the enhanced trimerization of RABV-G in complex with RVA122 likely results from RVA122 stabilizing the pre-fusion conformation by bridging domains I and III, rather than bridging protomers.

### Fusion loops stabilize soluble ectodomain trimers

In addition to RVA122 and the trimeric interface, RABV-G fusion loops also appear to play a role in trimerization and to influence protein stability. In a prior study, when RABV-G fusion loops were replaced with Gly-Ser linkers to facilitate protein purification and crystallization, no trimerization was observed at neutral or acidic pH (*7*). In contrast, we observe that wild-type RABV-G ectodomains containing intact fusion loops associate readily (Fig. S4). We also observe that when RABV-G trimers are not anchored to micelles or cellular membranes, the fusion loops are either disordered or interacting with each other (Fig. 1C, 4A). These results suggest that the fusion loops impact glycoprotein association and trimerization in addition to driving fusion of the viral and cell membranes after endosomal acidification.

**Fig. 4.**
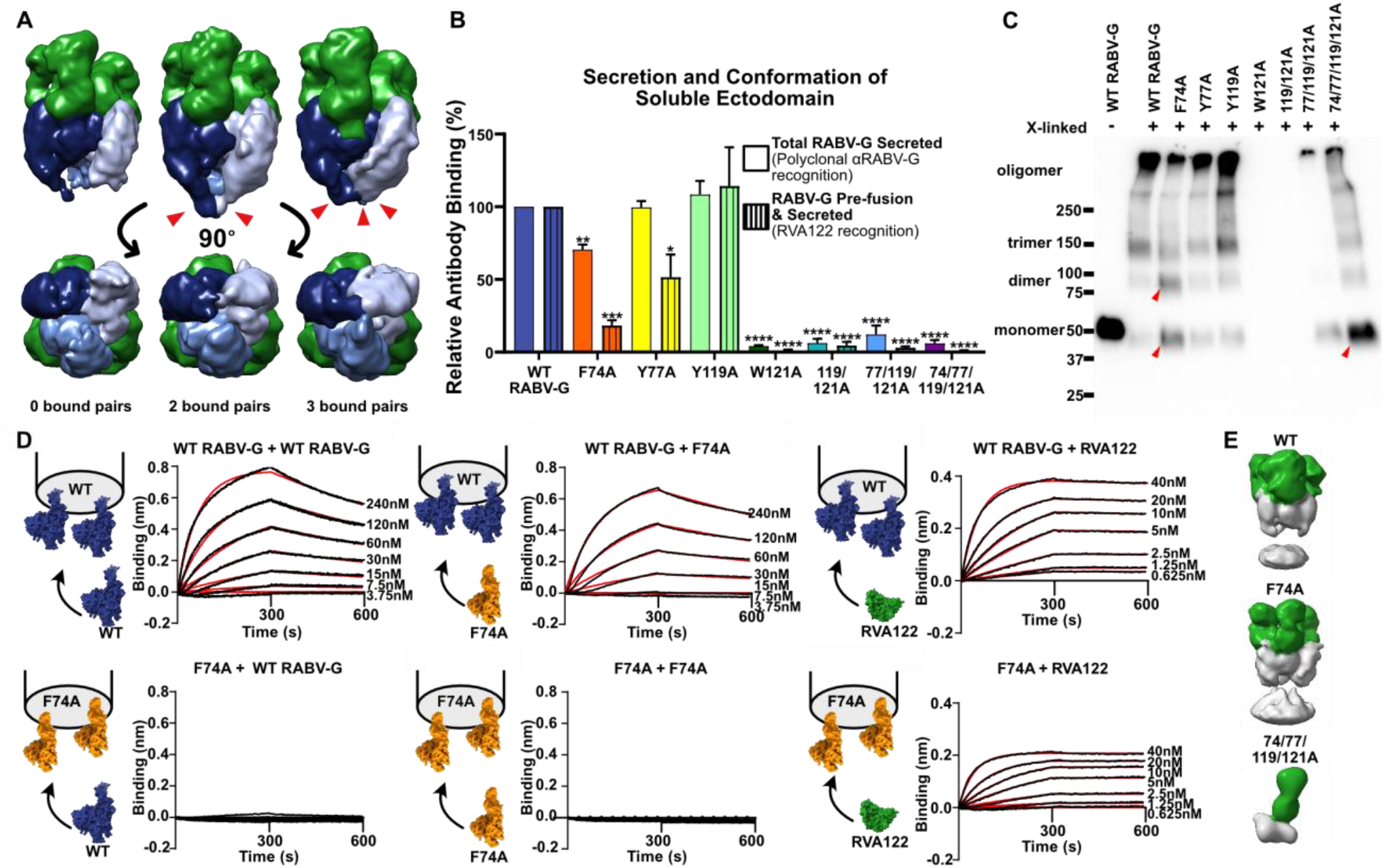
Fusion loop point mutations affect secretion, conformation, and trimerization of soluble RABV-G ectodomain. **(A)** Low-resolution cryo-EM reconstructions of RABV-G/RVA122 complexes with 0, 2, or 3 pairs (red arrowheads) of fusion loops interacting with each other. **(B)** ELISA quantifying secreted RABV-G and pre-fusion, secreted RABV-G for wild-type and mutant soluble ectodomains. **(C)** Crosslinking assay showing monomers, dimers, trimers, and higher-order oligomers formed from soluble RABV-G ectodomains. Red arrowheads indicate changes in oligomerization for F74A and 74/77/119/121A mutants compared to the wild-type. **(D)** Bio-layer interferometry experiments detailing binding interactions between wild-type RABV-G protomers, F74A RABV-G protomers, and RVA122 Fabs. **(E)** Negative stain EM reconstructions of soluble RABV-G (gray) immunoprecipitated with RVA122 Fabs (green). Statistical significance of differences between groups was analyzed using ANOVA. Error bars indicate SEM for three biological replicates. * p<0.05; ** p<0.01; ***p<0.001; ****p<0.0001.

To evaluate this hypothesis, we made alanine substitutions at aromatic fusion loop residues embedded in membranes and micelles (F74, Y77, Y119, and W121) both individually and in combination, and expressed these mutants as soluble ectodomains. We evaluated secretion and conformation of these mutants via ELISA, and oligomerization via western blot and negative stain electron microscopy. All fusion loop mutations except for Y119A significantly reduce the amount of secreted, pre-fusion RABV-G ectodomain compared to the wild-type (Fig. 4B). F74A results in a 30% reduction in total secreted RABV-G and 80% reduction in pre-fusion secreted RABV-G (Fig. 4B), whereas Y77A results in a 50% reduction of only secreted pre-fusion RABV-G. Mutation of W121 to alanine, alone or in combination with any other mutation, severely reduces or eliminates total glycoprotein secretion (Fig. 4B). Expression of W121A-containing constructs could be detected in cell lysates (Fig. S5), showing that the decreased secretion is not due to lower protein expression.

In addition to decreasing secretion of the soluble ectodomain, F74A also substantially reduces the amount of trimers, but not dimers, detected via protein crosslinking (Fig. 4C). In bio-layer interferometry experiments, F74A mutants bind RVA122 Fabs, indicating that some of the glycoprotein is in the pre-fusion conformation, but protomers do not stably associate with each other (Fig. 4D). F74A ectodomains immunoprecipitated and stabilized with RVA122, however, do trimerize (Fig. 4E). Interestingly, F74A protomers bind to wild-type RABV-G protomers in bio-layer interferometry experiments when wild-type RABV-G is the ligand (attached to the biosensor), but not the analyte (in solution) (Fig. 4D). These results suggest that F74A forms transient dimers in solution that can pair with wild-type protomers to form trimers and that RVA122-mediated stabilization of the pre-fusion conformation can also overcome the barrier to trimerization.

Soluble ectodomains containing the W121A mutation were poorly secreted and sufficient amounts of protein could not be produced for bio-layer interferometry experiments. W121A, 119/121A, and 77/119/121A were also too poorly secreted to be evaluated in crosslinking experiments. The quadruple mutant 74/77/119/121A forms primarily monomers in crosslinking experiments (Fig. 4C), and 74/77/119/121A soluble ectodomains immunoprecipitated with RVA122 also appear entirely monomeric via negative stain electron microscopy (Fig. 4E), indicating that any dimers or trimers that this mutant does form are unstable.

### Fusion loops stabilize full-length RABV-G pre-fusion conformation

In order to determine if the fusion loops also impact the conformation and stability of full-length RABV-G, we expressed the fusion loop point mutants as full-length glycoproteins, which include a transmembrane domain and cytoplasmic tail, and analyzed them with flow cytometry and immunofluorescence staining. All full-length versions of the RABV-G mutants express well and readily reach the cell surface (Fig. 5A). However, for mutants containing W121A, significantly less of the full-length protein is in the pre-fusion conformation (Fig. S6), and the pre-fusion RABV-G localizes to different areas of the cell compared to the wild-type (Fig. 5A).

In rhabdoviruses, the full-length glycoprotein can be either membrane-anchored or proteolytically cleaved near the transmembrane domain, releasing a shed ectodomain (*23*–*25*). Shed RABV-G is slightly longer than the cloned, soluble ectodomain (447 amino acids instead of 420) and contains no purification tags (*25*). The role of shed RABV-G during infection is unknown, but it may serve as a decoy for the immune system, as has been observed for other viruses (*26*–*28*). Furthermore, shed VSV-G has been reported to bind to full-length glycoprotein for uptake into cells (*23*) and injection of soluble RABV-G ectodomains into the brains of mice increases locomotion (*29*), a rabies symptom believed to facilitate transmission to new hosts.

**Fig. 5.**
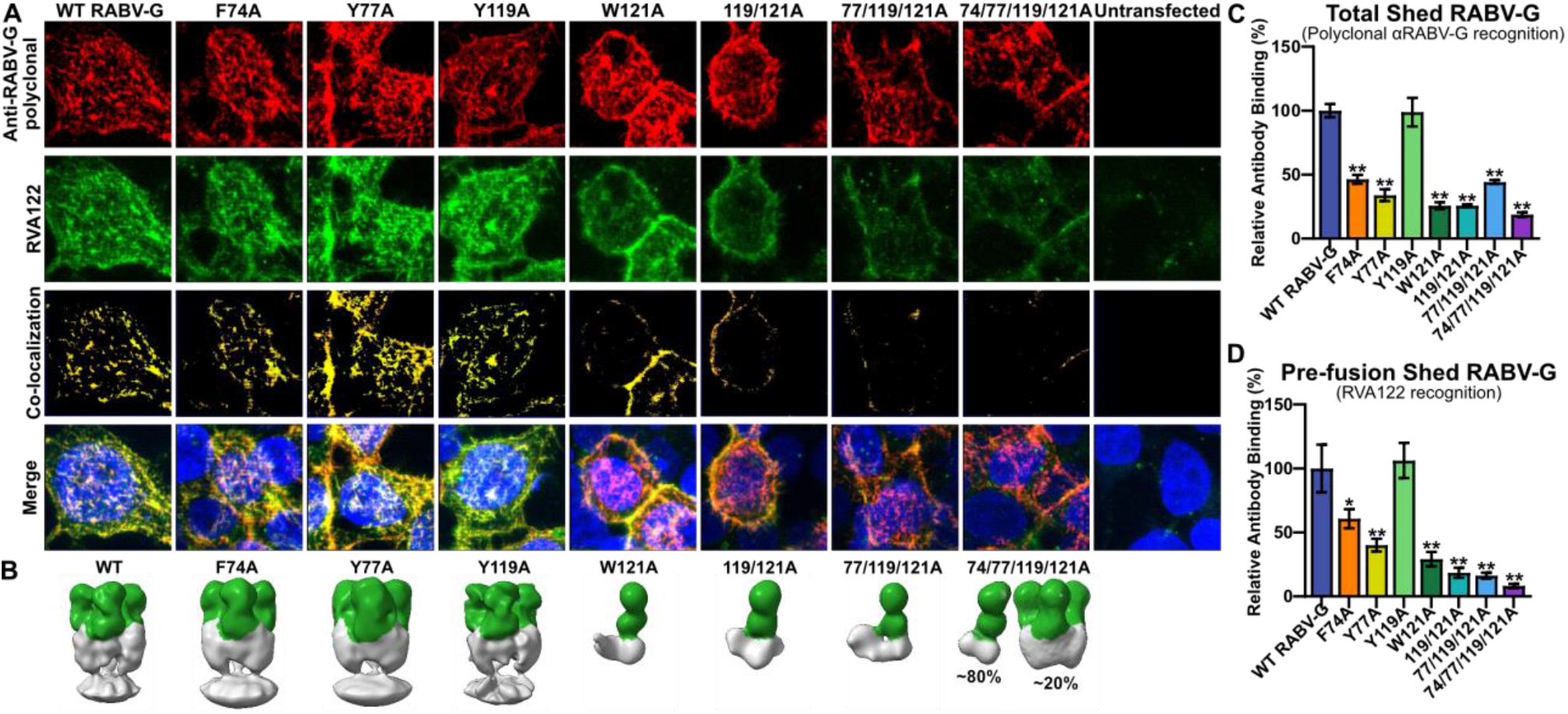
Fusion loop mutations impact full-length RABV-G conformation and trimerization. **(A)** Confocal immunofluorescence of cell surface-displayed, full-length RABV-G with fusion loop point mutations, stained for total RABV-G expression (polyclonal) and pre-fusion RABV-G (RVA122). Images are displayed as maximum intensity projections. **(B)** Negative stain electron microscopy reconstructions of shed RABV-G (gray) purified via immunoprecipitation with RVA122 (green). **(C-D)** ELISA quantification of the amount of total shed RABV-G (C) or pre-fusion shed RABV-G (D) carrying fusion loop mutations relative to the wild-type. Error bars indicate SEM for three biological replicates. * p<0.05; ** p<0.01.

We observe that fusion loop mutations impact the release of shed glycoprotein into tissue culture supernatant. All point mutations except Y119A decrease the total amount of shed RABV-G relative to the wild-type (Fig. 5C). Furthermore, there is also a significant decrease in the ratio of pre-fusion shed RABV-G to total shed RABV-G for the 77/119/121A and 74/77/119/121A mutants (Fig. S6). To visualize the structure of shed RABV-G, we immunoprecipitated mutants with RVA122 and examined them with negative stain electron microscopy. Shed wild-type, F74A, Y77A, and Y119A RABV-G each form trimers when in complex with RVA122, with the fusion loops embedded in cellular membranes that co-precipitate with the glycoprotein (Fig. 5B). In contrast, none of the mutants that contain W121A co-purify with cellular membranes. They are also mostly or entirely monomeric, with ∼20% of the RVA122-captured 74/77/119/121A RABV-G forming trimers and the other mutants containing W121A forming monomers (Fig. 5B).

Together, these results suggest that for the full-length glycoprotein, W121 stabilizes the pre-fusion conformation by interacting with membranes, preventing the glycoprotein from shifting into alternate conformations. The ability of shed 74/77/119/121A to trimerize in complex with RVA122, but not the other W121A mutants or 74/77/119/121A soluble ectodomain (Fig. 4E) is puzzling, but might be explained by relative protein expression levels or interactions among the fusion loops.

## Discussion

Rabies virus infection is over 99% lethal, still kills over 50,000 people annually, and is endemic in all populated continents. Rabies glycoprotein is a critical component of vaccines and target for antibody therapeutics and potential antiviral drugs. Despite considerable advances in the fields of immunology and vaccinology, rabies vaccines do not yet elicit lifelong immunity, leaving many individuals with no detectable rabies antibodies within one year of vaccination. Structural heterogeneity of the RABV-G likely contributes to the disappointing antibody response. Pre-fusion, trimeric RABV-G would be an ideal immunogen for vaccination, but presentation of this form has been challenging due to the inherent instability of the molecule. Lack of information on what the trimeric structure is, how it is organized, and how it can be stabilized has hampered these efforts.

Here we report the structure of trimeric, pre-fusion RABV-G bound to a potent and broadly neutralizing antibody. In this structure, the fusion loops are anchored into cellular membrane/micelle density, a feature not appreciated in a previously published structure of the pre-fusion RABV-G monomer, in which the fusion loops were replaced by linkers. We show that the fusion loops play an integral role in stabilizing the pre-fusion conformation, regardless of whether RABV-G is full-length, shed, or expressed as a soluble ectodomain. Hence, inclusion of these elements in vaccine designs will be critical for the faithful display of the pre-fusion RABV-G trimer. A nucleic acid-based vaccine may bypass issues associated with purification, but must still direct expression of glycoproteins with the correct conformation and quaternary assembly. The pre-fusion, trimeric RABV-G structure illustrated here, the demonstration of the importance of fusion loops for successful trimerization and conformational stabilization, and the visualization of the potent and broadly neutralizing antibody epitope, together provide a key road map for development of improved vaccines and post-exposure therapeutics to prevent the deaths from rabies infection.

## Acknowledgments

We thank Sharon Schendel for manuscript editing.

## Author contributions

Conceptualization: HMC, EOS

Formal Analysis: HMC, DZ

Funding acquisition: EOS, HMC, DZ

Investigation: HMC, RD, AA

Methodology: HMC, DZ

Resources: DC, HB

Validation: HMC, KH, DZ

Visualization: HMC

Writing – original draft: HMC

Writing – review & editing: HMC, KH, EOS

## Data and materials availability

The cryo-EM density map was deposited in the Electron Microscopy Data Bank under the EMDB accession code XXX and the atomic model was deposited in the Protein Data Bank under the PDB accession code YYY.

## Supplementary Materials

### Materials and Methods

#### Cells

Antigen binding fragments (Fabs) were prepared in S2 insect cells (*Drosophila melanogaster*, Invitrogen #R69007) grown in Insect-XPRESS media (Lonza) with 1% penicillin-streptomycin (Thermofisher) at 27 °C in a rotary shaker. RABV-G ectodomain and full-length protein was produced in HEK-293T cells (*Homo sapiens*, ATCC #CRL-3216, RRID:CVCL_0063) grown in T75 flasks (Corning) in DMEM (Thermofisher) with 10% fetal bovine serum (FBS) (Gibco) and 1% penicillin-streptomycin (Thermofisher) at 37 °C with 5% CO_2_.

#### Plasmids and cloning

Codon-optimized sequences for the heavy and light variable domains of the neutralizing antibody RVA122 were synthesized and cloned into pMT puro and pMT plasmids, respectively, containing the corresponding regions for Fab IgG1 heavy and IgK light chains. The heavy chain sequence also contained a C-terminal enterokinase cleavage site, followed by a double-strep tag.

The codon-optimized sequence for full-length, PV strain RABV-G (NCBI reference sequence: NC_001542.1) was synthesized and cloned into the pcDNA3.1(-) vector. The soluble ectodomain was cloned by truncating the full-length glycoprotein to residues 1-439 (numbering includes signal peptide) and adding either no tag, a V5/6x-His tag, or a double strep-tag and an avi-tag to the C-terminus. Point mutations to full length and soluble ectodomain plasmids were made via site-directed mutagenesis using Q5 DNA polymerase (NEB), T4 polynucleotide kinase (NEB), and T4 DNA ligase (NEB).

#### Protein expression and purification

For RVA122 expression, heavy chain and light chain plasmids were co-transfected into S2 cells using Effectene (Qiagen) according to the manufacturer’s protocol, and selected using puromycin (Invivogen). Cells were expanded after selection, transferred to shaker flasks (TriForest Enterprises), and induced with CuSO_4_ (500*μ*M) and sodium butyrate (5mM) once cells had reached a density of ∼1×10^7^ cells/cm^2^. Cells were harvested 4 days post induction and pelleted at 4,000g. Supernatant was saved and adjusted to pH 8.0 with NaOH, then freeze-thawed at – 20°C/25°C to precipitate salts from the supernatant prior to purification. Supernatant was filtered and Fabs were purified on a StrepTrap HP column (GE Healthcare). Eluted fractions were pooled and concentrated using a 10 kDa MWCO centrifugal filter (Millipore), followed by further fractionation over an S200i column (GE Healthcare) in PBS at 0.5 mL/min to remove aggregates.

For RABV-G ectodomain expression, HEK 293T cells seeded in T75 flasks (Corning) at a density of 4×10^4^ cells/cm^2^ were transiently transfected with 9.8μg of the recombinant plasmid using Polyethyleneimine (PEI) at a 3:1 PEI:DNA ratio. At two and four days post transfection, supernatant was collected from flasks and centrifuged at 4,000g to remove cell debris prior to purification. Tagged RABV-G ectodomain was purified from supernatant with NiNTA agarose beads (Qiagen) for His-tagged protein or Streptactin Superflow plus beads (Qiagen) for strep-tagged protein. Untagged RABV-G ectodomain was purified from supernatant via immunoprecipitation with the strep-tagged RVA122 Fab on Streptactin Superflow plus beads (Qiagen).

Briefly, the supernatant was adjusted to pH 8.0 with NaOH, then incubated overnight at 4°C with either beads for tagged RABV-G ectodomain, or with RVA122 Fab-captured beads. Beads were pelleted from supernatant via centrifugation, washed 3 times with PBS, then eluted multiple times with either 250mM imidazole/TBS for NiNTA beads or 10mM d-desthiobiotin/PBS for Streptactin beads. Protein for cryo-electron microscopy was concentrated and buffer exchanged into PBS in a 100kDa MWCO centrifugal filter (Millipore), which removed unbound RVA122 Fabs.

Shed RABV-G was produced by expression of full-length RABV-G in 293T cells and collection of supernatant four days after transfection. Purified, shed RABV-G was produced via immunoprecipitation with RVA122 Fabs, as described above.

#### Preparation of cryoEM grids and data collection

Purified RABV-G/RVA122 complexes at a concentration of ∼200 μg/mL were mixed 3:1 with 0.12mM (0.03mM final concentration) Lauryl Maltose Neopentyl Glycol (LMNG) (Anatrace) and immediately frozen on 1.2/1.3 μm C-flat grids (EMS) using an FEI vitrobot Mark IV (ThermoFisher) in 85% humidity and 4°C with a 10s blot time. A total of 1,969 micrographs were collected on a 200keV Talos Arctica with a K2 direct electron detector (Gatan) at the Pacific Northwest Cryo-EM Center (PNCC) with pixel size 1.15 Å/pixel and a total dose of 26.7 e^-^/Å^2^.

A second cryo-EM dataset of 6,305 micrographs was collected with ∼200 μg/mL purified RABV-G/RVA122 complexes mixed 3:1 with 0.36mM (0.09mM final concentration) LMNG and frozen on 2/1 μm C-flat grids in 85% humidity and 4°C with a 10s blot time. Images were collected on a 300keV Titan Krios electron microscope with a K3 direct electron detector at 1.1 Å/pixel and a total dose of 50.0 e^-^/Å^2^.

#### CryoEM data processing and model building

The dataset containing RVA122/RABV-G complexes with 0.09mM LMNG (Table S1) was used for high resolution reconstruction and model building. Recorded movies were motion corrected and dose weighed using either cryoSPARC’s own algorithm or RELION’s MotionCorr2 implementation. CTF parameters were determined either with cryoSPARC’s Patch CTF program or in RELION using CTFFIND4 (*30*).

Particles were picked using the TOPAZ neural network picker (*31*) and processed in cryoSPARC (*32*). 2D classification and 3D hetero-refinement were used to screen for particles bound to cellular membrane or micelle densities. Symmetry expanded particles were used in a local refinement job to produce the first high-resolution map. To further improve the map quality around the fusion loops, particle coordinates were transferred to RELION 3.1 (*33*). In RELION, particles were sorted via 3D classification without alignment to isolate complexes with better resolved fusion loops. Per-particle CTF parameters were estimated and particles were Bayesian polished to yield the final map. Final half-maps were transferred to cryoSPARC to estimate the local resolution. A detailed flowchart containing data processing steps is presented in Figure s7.

Model building was performed in COOT 0.9.2 (*34*) and ISOLDE (*35*). Model refinement and validation was performed in Phenix 1.19 (*36*). Model building was aided by a protein model generated from AlphaFold2 (*37, 38*). ChimeraX 1.2.5 (*39*) was used to prepare figures of the structure. Fourier shell correlation curves were generated using RELION, the local resolution histogram using cryoSPARC, and correlation coefficient graphs using Phenix.

Lower resolution cryo-EM maps of RABV-G/RVA122 complexes not bound to micelles or cellular membranes were prepared from the dataset containing complexes with 0.03mM LMNG. Particles were picked with TOPAZ, as above, reconstructed into an initial model in cryoSPARC with C3 symmetry, and then sorted via 3D variability analysis focusing on the fusion loops in order to identify particles with fusion loops in different conformations. Groups of similar particles were reconstructed with a homogeneous refinement job and C1 symmetry.

#### Negative Stain Electron Microscopy and Reconstruction

Shed RABV-G and soluble RABV-G were purified via immunoprecipitation with RVA122 Fabs, as described above, then added to C-flat CF400Cu grids (EMS) at a concentration of approximately 10*μ*g/mL and stained with 0.5% uranyl acetate. Grids were imaged on a Titan Halo electron microscope operated at 300keV with a Falcon 3 direct electron detector at 1.87 pixels/Å and 50 e^-^/Å^2^. Particles were picked with TOPAZ and reconstructed in Cryosparc using homogeneous refinement and C3 symmetry.

#### ELISAs

HEK293T cells were seeded in 6 well plates (Corning) at a density of 4×10^4^ cells/cm^2^ and grown overnight. Cells were then transfected with plasmids encoding full length or soluble rabies glycoprotein using PEI at a ratio of 1:3 DNA to PEI (1.26*μ*g DNA and 3.8*μ*g PEI per well). Four days post-transfection, the supernatant was removed from cells, diluted 1:1 in PBS, then transferred to half-well ELISA plates (Corning) and incubated for 1hr at room temperature. The supernatant was removed and wells were blocked with 3% bovine serum albumin (BSA) (Sigma-Aldrich) in PBS for an additional hour at room temperature. Wells were washed once with PBS with 0.05% CHAPS (3-[(3-cholamidopropyl)dimethylammonio]-1-propanesulfonate) (BioVision), then incubated with either 1:2,000 rabbit anti-RABV-G polyclonal antibody (a kind gift of Dr. Mattias Schnell, Thomas Jefferson University) or RVA122 monoclonal antibody for 1hr at room temperature. Plates were washed three times with PBS/0.05% CHAPS, then incubated with either 1:2,000 goat anti-human Fab HRP (Jackson ImmunoResearch Labs Cat# 109-036-006, RRID:AB_2337590) or 1:2,000 goat anti-rabbit HRP (SouthernBiotech Cat# 4050-05, RRID:AB_2795955) for 1hr at room temperature. Plates were washed three times with PBS/0.05% CHAPS, then developed with 1-step Ultra TMB-ELISA substrate solution (Fisher) and quenched with 1M sulfuric acid (Fisher). Plates were read at 450 nm on a Tecan Spark 10M plate reader. Three biological replicates were performed for each experiment and results were analyzed via ANOVA in Graphpad Prism 9.

#### Protein crosslinking assays

Purified His-tagged RABV-G ectodomains were buffer exchanged into PBS, diluted to a concentration of 56μg/mL, and incubated in 0.01% glutaraldehyde/PBS for 1hr at room temperature. Crosslinking was quenched via addition of 1M Tris pH 8.0 to a final concentration of 50mM. Proteins were separated via SDS-PAGE gel under reducing conditions, transferred to a PVDF membrane (Millipore), and stained with 1:2,000 rabbit polyclonal anti-RABV-G antibody and 1:2,000 goat anti-rabbit HRP antibody (SouthernBiotech Cat# 4050-05, RRID:AB_2795955) for visualization.

#### Measurement of mutant RABV-G ectodomain expression levels

293T cells seeded in 6-well plates at a density of 4×10^4^ cells/cm^2^ were transfected with mutant RABV-G ectodomains as described above. Four days post transfection, cells were washed with PBS, lysed with 1% Triton X-100 in PBS, separated via SDS-PAGE gel under reducing conditions, transferred to a PVDF membrane, and stained with 1:2,000 rabbit polyclonal anti-RABV-G antibody and 1:2,000 mouse anti-βactin antibody (Santa Cruz Biotechnology #sc-69879, RRID:AB_1119529), then 1:2,000 goat anti-mouse HRP antibody (Thermo Fisher Scientific Cat# 31437, RRID:AB_228295) and 1:2,000 goat anti-rabbit HRP antibody (SouthernBiotech Cat# 4050-05, RRID:AB_2795955) for visualization.

#### Immunofluorescence Assays and Flow Cytometry

HEK293T cells were seeded on 12mm circular glass coverslips (Fisher) in a 24 well plate (Corning) at a density of 4×10^4^ cells/cm^2^ and grown overnight. Cells were then transfected with plasmids encoding full length RABV-G using PEI at a ratio of 1:3 DNA to PEI (0.25*μ*g DNA and 0.75*μ*g PEI per well). Two days post transfection, cells were fixed in 4% paraformaldehyde/PBS for 10 min at room temperature, then stained with 1:2,000 rabbit polyclonal anti-RABV-G and 1:2,000 RVA122 monoclonal antibody in PBS with 0.1% BSA for 1hr at room temperature. Cells were washed twice with PBS, then stained with 1:2,000 anti-human FITC (SouthernBiotech Cat# 2040-02, RRID:AB_2795641) and 1:2,000 anti-rabbit Alexa 568 (Thermo Fisher Scientific Cat# A-11011, RRID:AB_143157) for 1hr at room temperature. Cells were washed twice with PBS, then stained with 1:10,000 10mg/mL Hoechst for 15 min and washed with PBS. Coverslips were mounted on slides with Prolong Gold Antifade Mountant (ThermoFisher), and imaged on a Zeiss 880 confocal microscope at 40x magnification.

For flow cytometry experiments, HEK293T cells were similarly seeded in 6-well plates and transfected with plasmids encoding full-length RABV-G. Two days post-transfection, cells were fixed with 4% PFA, washed with PBS, incubated in Human BD Fc Block (BD Biosciences) and stained with 1:1,000 anti-RABV-G polyclonal antibody and 1:1,000 RVA122 monoclonal antibody for 1hr. Cells were washed twice, then stained with 1:1,000 goat anti-rabbit Alexa 647 (Thermo Fisher Scientific Cat# A27040, RRID:AB_2536101) and 1:1,000 goat anti-human FITC (SouthernBiotech Cat# 2040-02, RRID:AB_2795641) for 1hr. Cells were washed twice, then quantified using a LSR-II flow cytometer (BD Biosciences). Experiments were performed in triplicate (biological replicates), with untransfected cells serving as a negative control. RVA122 mean fluorescence intensity (MFI) was analyzed via ANOVA in Graphpad Prism 9.

#### Bio-layer Interferometry (BLI)

RABV-G ectodomains used to quantify binding kinetics contained a C-terminal double strep tag and avi-tag. RABV-G used as a ligand was biotinylated at the avi-tag using BirA ligase according to the manufacturer’s protocol (Avidity), then buffer exchanged into PBS to remove unlinked biotin. Binding kinetics were measured using an Octet Red 384 (Fortebio). Streptavidin biosensors (Fortebio) were hydrated in kinetics buffer (PBS pH 7.4 with 0.1% BSA and 0.02% CHAPS) for 10 minutes at room temperature. Baseline readings were collected in kinetics buffer for 30s at 30°C, followed by loading of biotinylated RABV-G onto biosensors at a concentration of ∼3.5ug/mL for 300s and a 60s wash in kinetics buffer. For measurements of Fab binding affinity, RVA122 wild-type and mutant Fabs at concentrations ranging from 1.25-80nM were incubated with biosensors for 300s to measure association, followed by a 300s incubation in kinetics buffer to measure disassociation. For measurements of RABV-G binding affinity, biosensors were incubated with unbiotinylated RABV-G ectodomain at concentrations ranging from 3.75-960nM, followed by a 300s incubation in kinetics buffer. For measurements at pH 5.5, all steps were performed in 0.1M citric acid/trisodium citrate buffer pH 5.5 with 0.1% BSA and 0.02% CHAPS. Experiments were performed in duplicate (technical replicates) and a sensor run without analyte was used as a reference. Affinity rate constants were calculated by subtracting the reference, then applying global fitting using the Octet Data Analysis HT software version 11 (Fortebio).

**Fig. S1.**
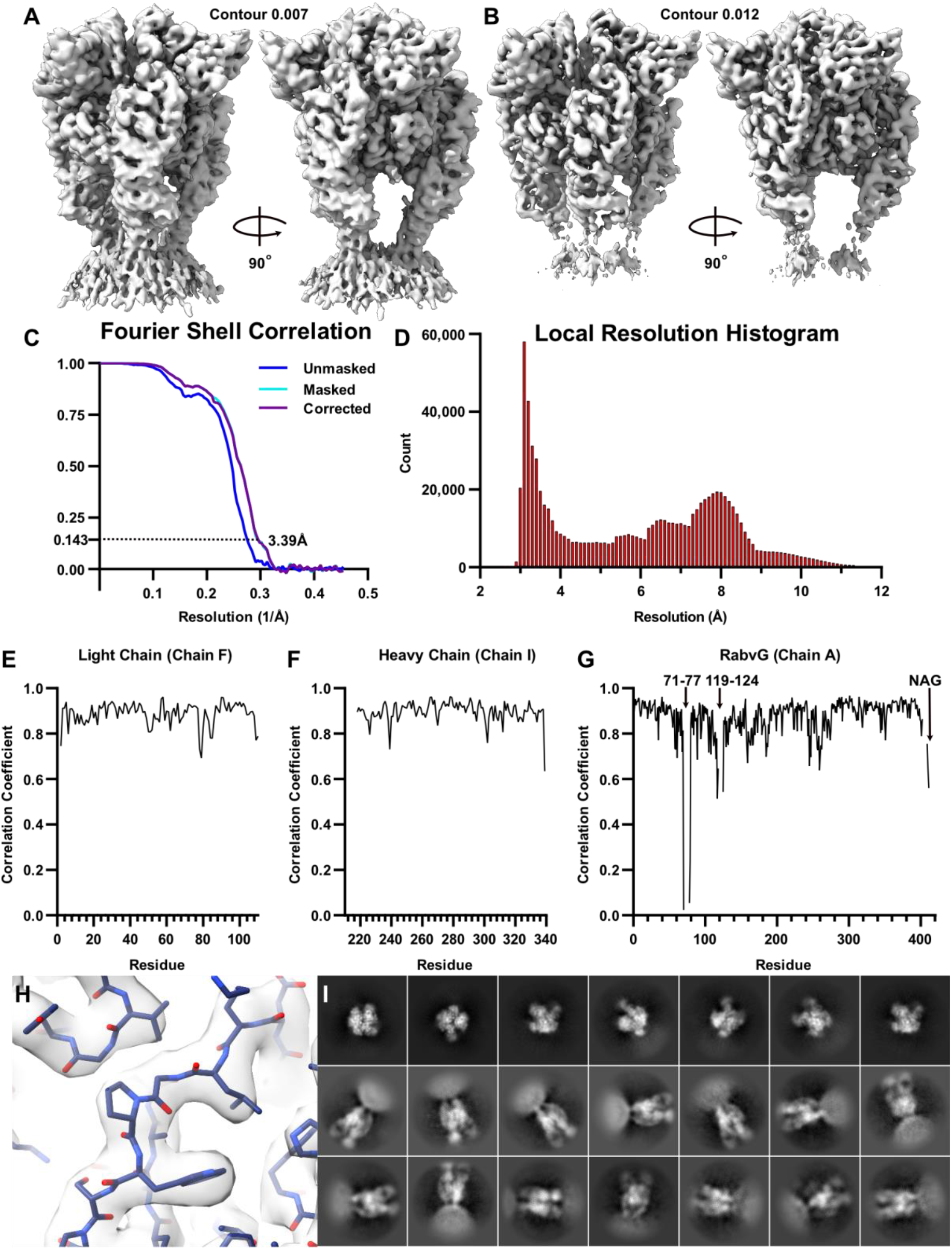
Additional information and statistics for the cryo-EM structure of RABV-G complexed with neutralizing antibody RVA122. **(A-B)** Electron density maps at low (A) and high (B) contour levels. **(C)** Fourier Shell correlation curve. **(D)** Local resolution histogram. **(E-G)** Correlation coefficient graphs for the RVA122 light chain (E), RVA122 heavy chain (F), and RABV-G (G). **(H)** Zoomed map showing the atomic model fit into electron density. **(I)** Representative 2D classes of cryo-EM particles used in 3D reconstruction.

**Fig. S2.**
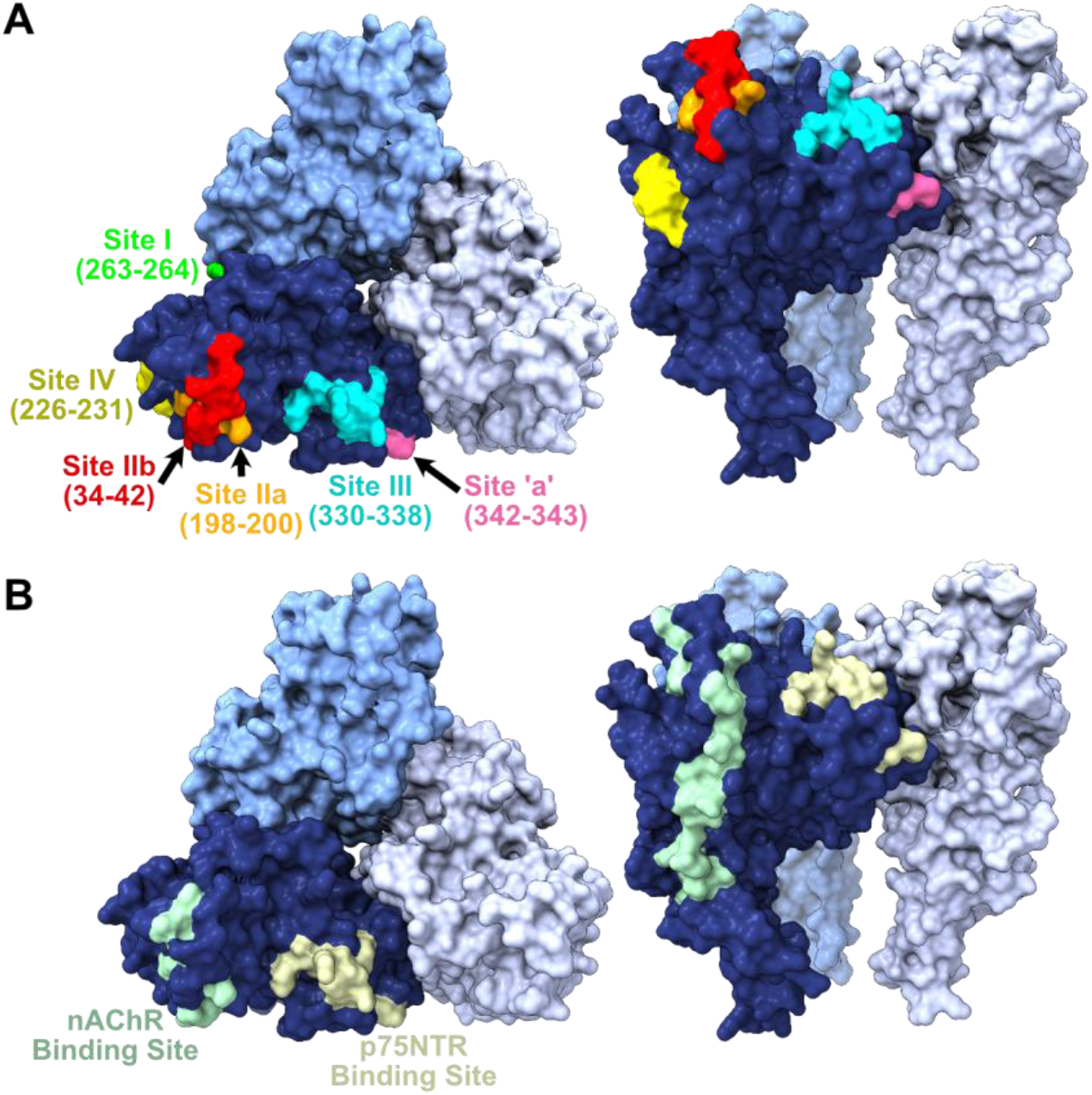
RABV-G antigenic sites and receptor binding sites. **(A)** Known RABV-G antigenic sites mapped onto the trimeric RABV-G structure, with residue numbers indicated. **(B)** Approximate receptor binding sites for the nicotinic acetylcholine receptor (nAChR) and the p75 neurotrophin receptor (p75NTR).

**Fig. S3.**
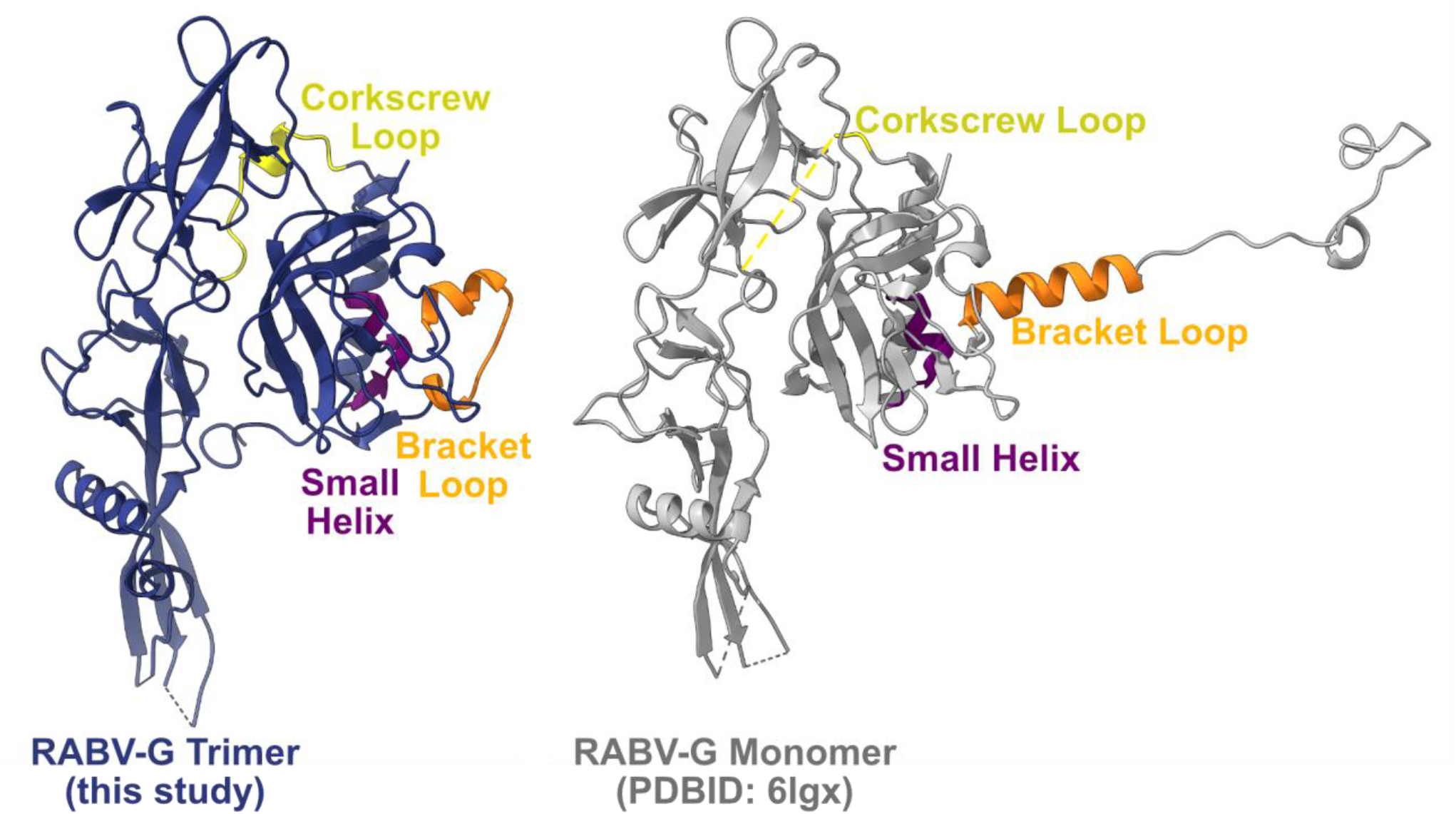
Pre-fusion RABV-G trimers and alternate pre-fusion RABV-G monomers have different conformations of loops at the trimeric interface. Structure of one RABV-G protomer from this study (left) and a RABV-G monomer from a crystal structure (PDBID: 6lgx) (right), with the small helix (purple), corkscrew loop (yellow), and bracket loop (orange) marked.

**Fig. S4.**
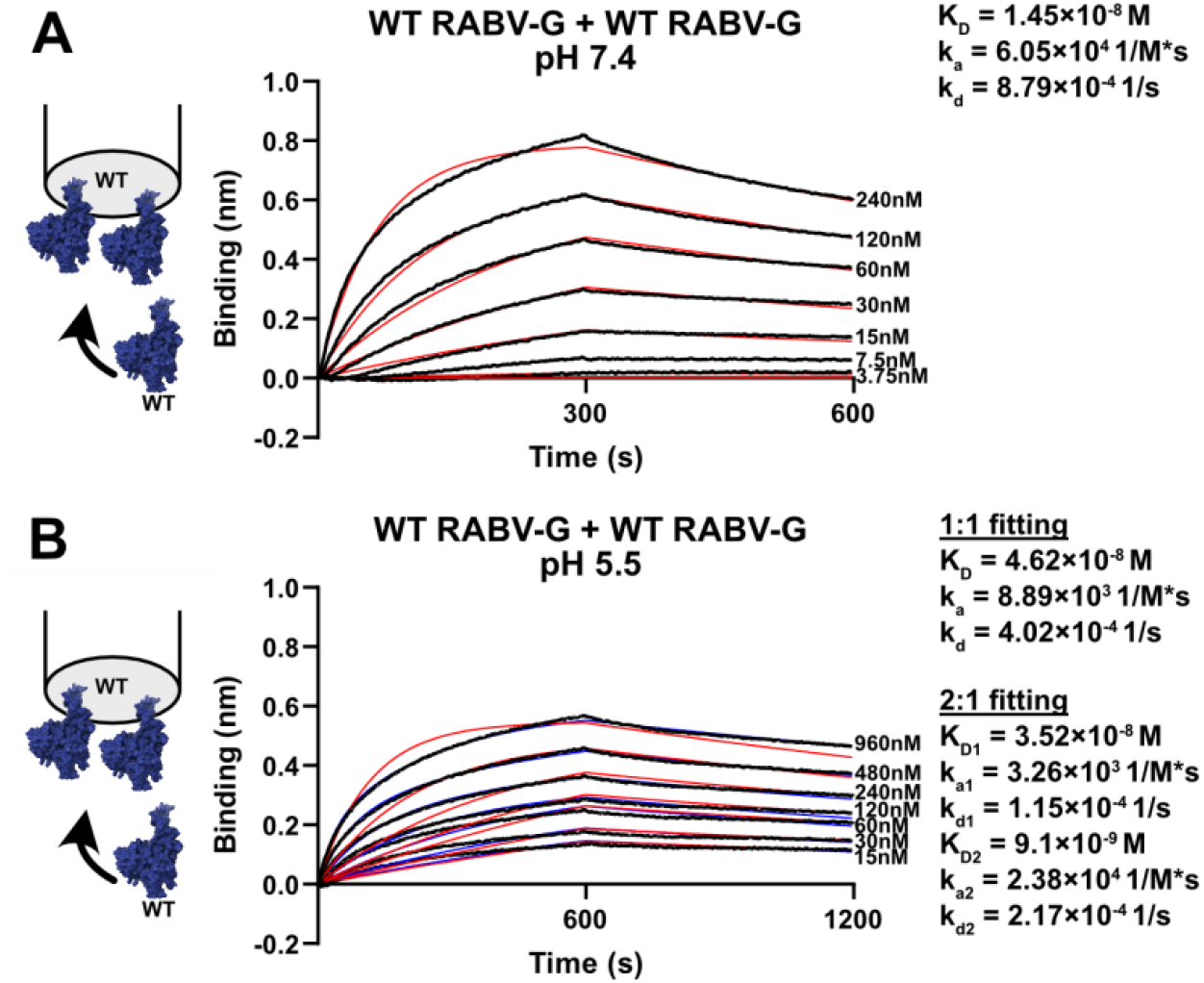
RABV-G soluble ectodomain binding kinetics. Bio-layer interferometry measurements of RABV-G protomer binding at pH 7.4 **(A)** and pH 5.5 **(B)**. 1:1 curve fitting (red, pH 7.4 and 5.5) and 2:1 curve fitting (blue, pH 5.5 only) are shown.

**Fig. S5.**
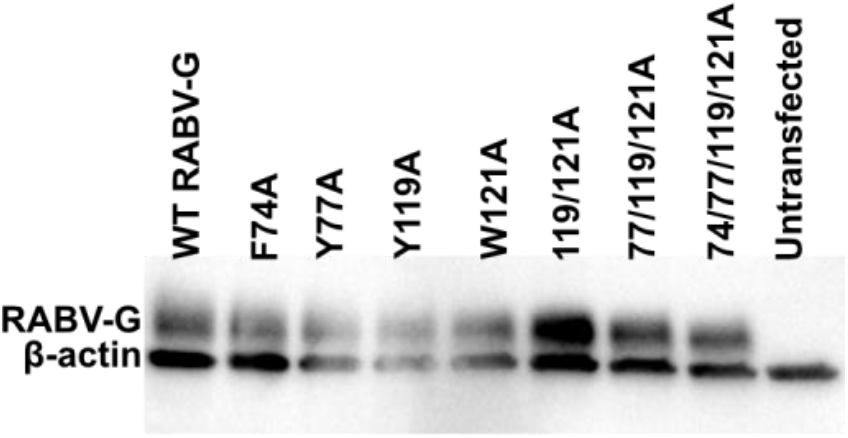
Expression of RABV-G soluble ectodomains. Western blot showing total protein expression levels of RABV-G soluble ectodomains and β-actin (loading control) from lysed cells.

**Fig. S6.**
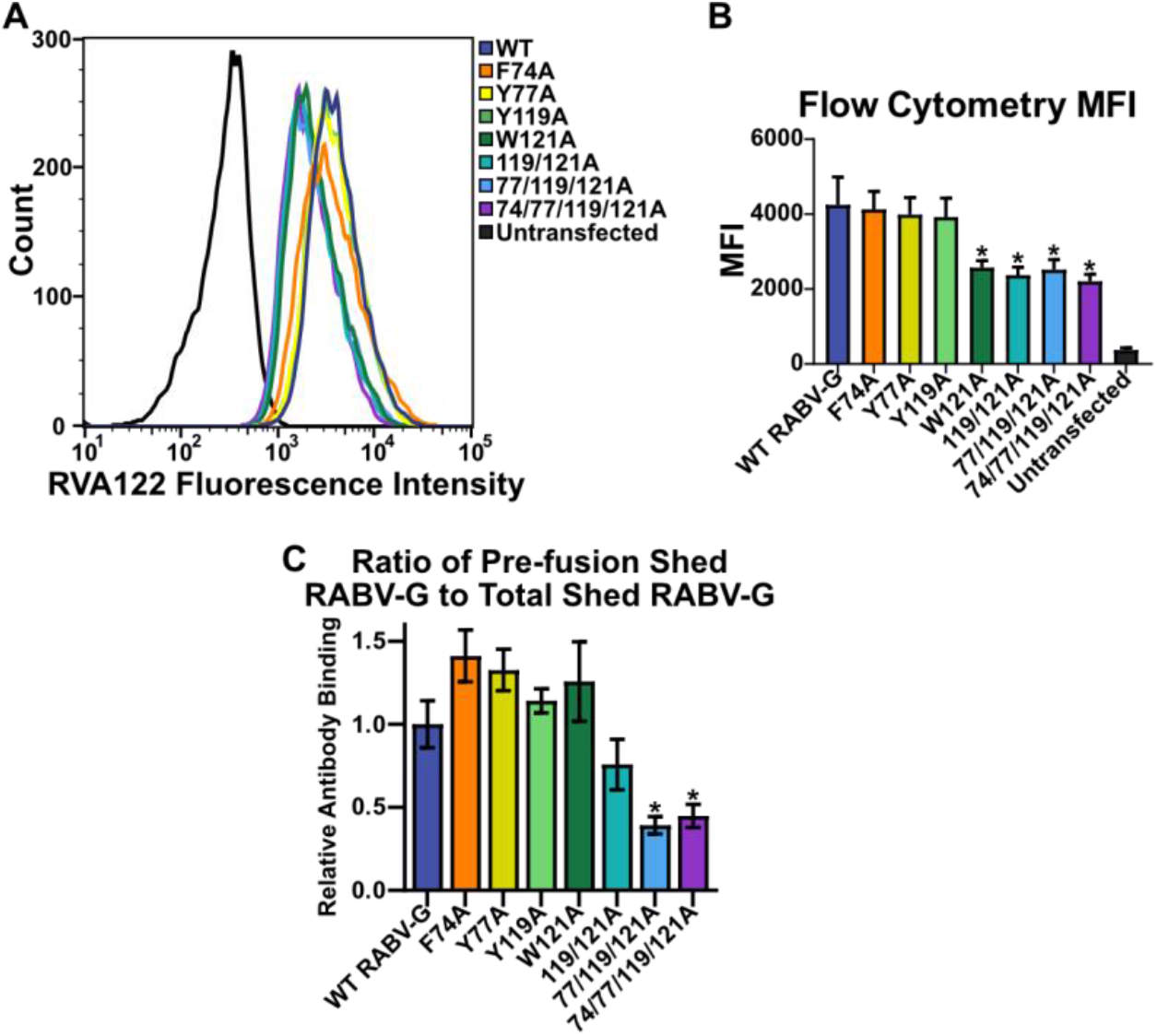
Fusion loop point mutants affect protein conformation and shedding for full-length RABV-G. **(A-B)** Flow cytometry quantifying the level of surface-expressed, full-length RABV-G in the pre-fusion conformation. **(C)** Ratio of pre-fusion shed RABV-G to total shed RABV-G in an ELISA quantifying shed RABV-G. Statistical significance of differences were analyzed using ANOVA. Error bars indicate SEM for three biological replicates. * p<0.05; ** p<0.01.

**Fig. S7.**
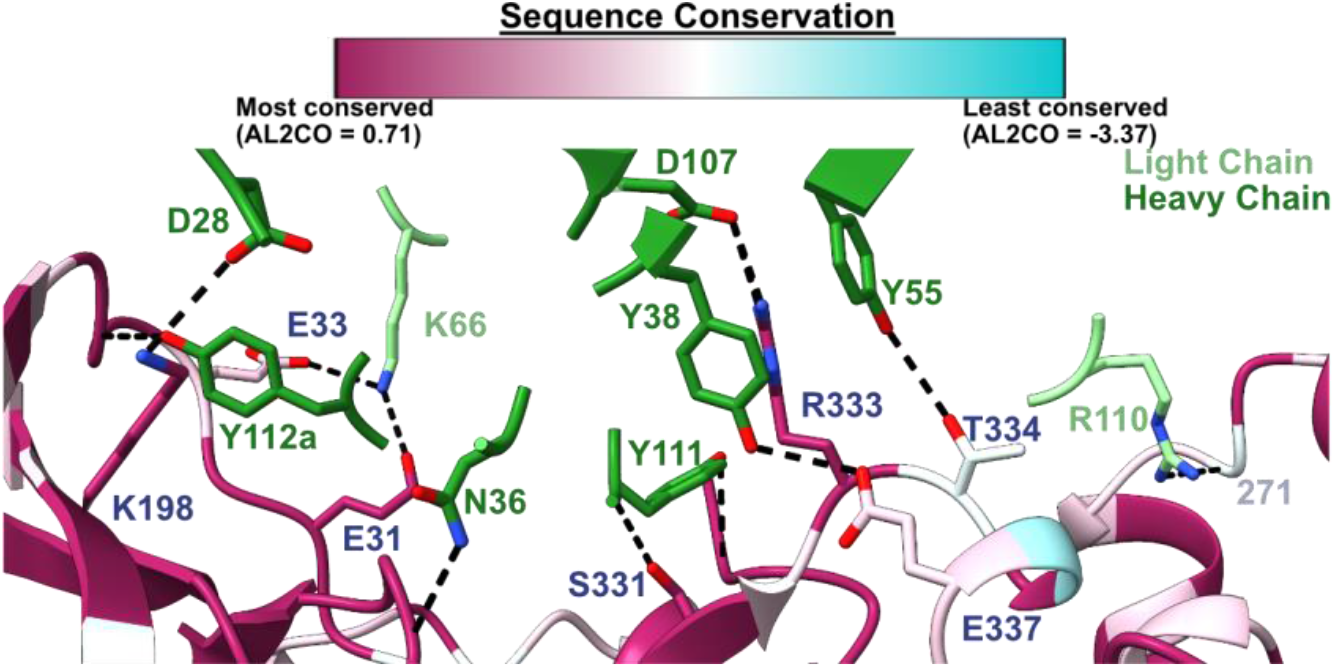
Sequence conservation among type 1 lyssaviruses at the RVA122 binding site. A sequence alignment was created from European Bat Lyssaviruses 1 (YP_001285391.1) and 2 (YP_001285396.1), Khujand virus (YP_009094330.1), Aravan Lyssavirus (YP_007641395.1), Bokeloh Bat Lyssavirus (YP_009091812.1), Duvenhage virus (NC_020810.1), Irkut virus (AFP74571.1), and PV strain rabies virus (NC_001542.1). ChimeraX 1.2.5 (*39*) was used to color RABV-G residues according to sequence conservation.

**Fig. S8.**
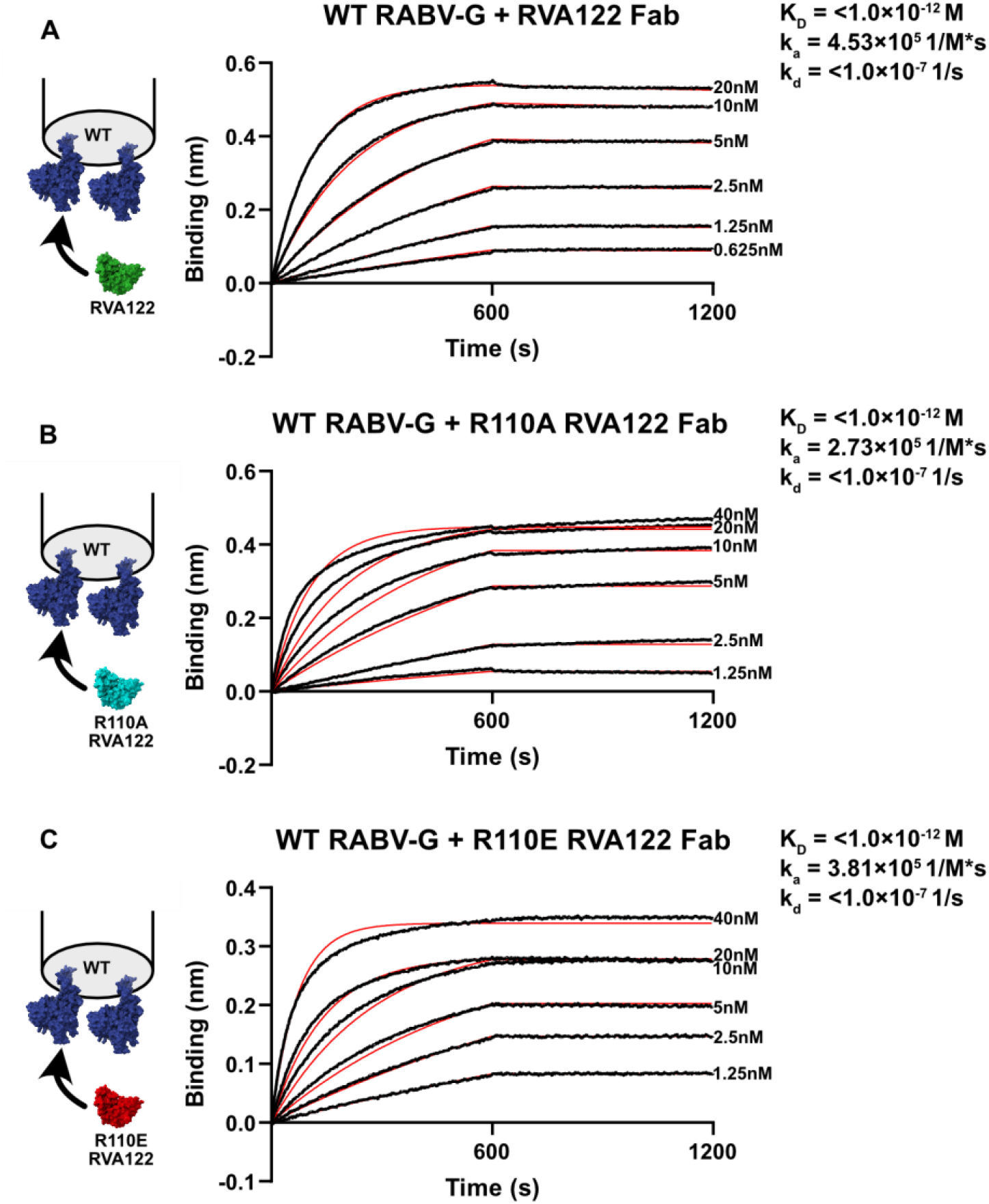
Point mutations to RVA122 residue R110 do not impact RABV-G binding affinity. Bio-layer interferometry measurements of binding kinetics between RABV-G and wild-type RVA122 Fab **(A)**, R110A RVA122 Fab **(B)**, and R110E RVA122 Fab **(C)**. 1:1 curve fitting (red) is shown.

**Fig. S9. Supplemental Figure 9.**
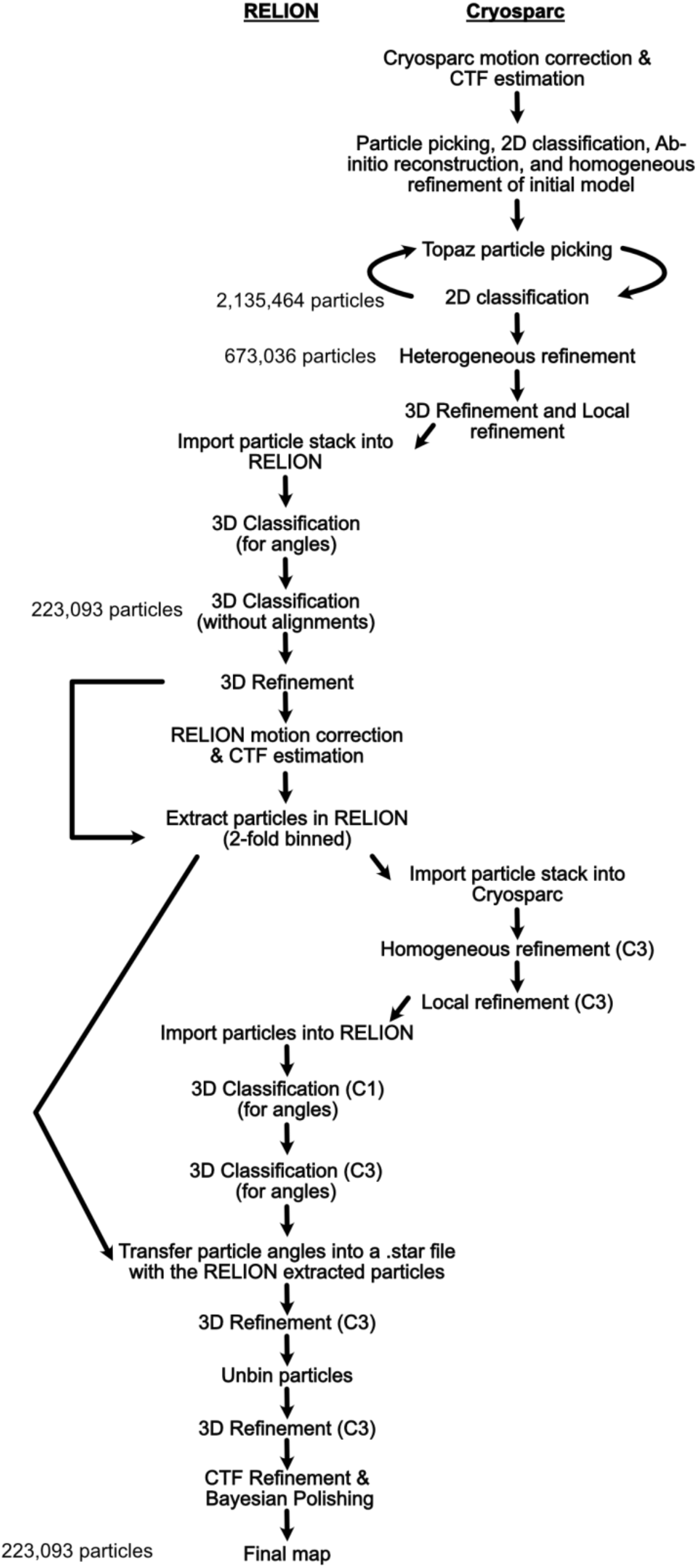
Cryo-EM data processing method for the high-resolution RABV-G/RVA122 complex.

**Table S1.** CryoEM map and model statistics.

|  | High Resolution | Low Resolution |  |  |
| --- | --- | --- | --- | --- |
|  | RABV-G/RVA122 Complex | RABV-G/RVA122 Complex |  |  |
|  |  | 3 engaged fusion loops | 2 engaged fusion loops | 0 engaged fusion loops |
| **Map Statistics** |  |  |  |  |
| Movies | 6,305 | 1,969 |  |  |
| Particles | 223,092 | 9,627 | 32,409 | 10,144 |
| B-factor | 140.5 | 299.9 | 256.7 | 382.7 |
| Resolution (Å) | 3.39 | 7.05 | 5.71 | 6.92 |
| Box size (px) | 256 | 256 |  |  |
| Pixel size (Å/px) | 1.1 | 1.15 |  |  |
| Microscope | Titan Krios | Talos Arctica |  |  |
| Detector | Gatan K3 | Gatan K2 |  |  |
| Total dose (e<sup>-</sup>/Å<sup>2</sup>) | 50 | 26.74 |  |  |
| Frames | 50 | 55 |  |  |
| Dose per frame (e<sup>-</sup>/Å<sup>2</sup>) | 1 | 0.49 |  |  |
| **Molprobrity statistics** |  |  |  |  |
| All-atom clashscore | 5.16 |  |  |  |
| Ramachandran plot: |  |  |  |  |
| Outliers | 0.00% |  |  |  |
| Allowed | 3.11% |  |  |  |
| Favored | 96.89% |  |  |  |
| Cbeta outliers | 0.00% |  |  |  |
| Rotamer outliers | 0.00% |  |  |  |
| Peptide plane: |  |  |  |  |
| Cis-proline | 0.00% |  |  |  |
| Cis-general | 0.00% |  |  |  |
| Twisted proline | 0.00% |  |  |  |
| Twisted general | 0.00% |  |  |  |

**Movie S1. 3D variability analysis of RABV-G fusion loops**.

